# Proteins linked to autosomal dominant and autosomal recessive disorders harbor characteristic rare missense mutation distribution patterns

**DOI:** 10.1101/018648

**Authors:** Tychele N. Turner, Christopher Douville, Dewey Kim, Peter D. Stenson, David N. Cooper, Aravinda Chakravarti, Rachel Karchin

**Author notes:** Send all correspondence to: Rachel Karchin, Ph.D., Departments of Biomedical Engineering and Oncology, Institute of Computational Medicine, Johns Hopkins University, Hackerman Hall 217A, Baltimore, MD 21210, E.

## Abstract

The role of rare missense variants in disease causation remains difficult to interpret. We explore whether the clustering pattern of rare missense variants (MAF<0.01) in a protein is associated with mode of inheritance. Mutations in genes associated with autosomal dominant (AD) conditions are known to result in either loss or gain of function, whereas mutations in genes associated with autosomal recessive (AR) conditions invariably result in loss of function. Loss-of-function mutations tend to be distributed uniformly along protein sequence, while gain-of-function mutations tend to localize to key regions. It has not previously been ascertained whether these patterns hold in general for rare missense mutations. We consider the extent to which rare missense variants are located within annotated protein domains and whether they form clusters, using a new unbiased method called CLUstering by Mutation Position (CLUMP). These approaches quantified a significant difference in clustering between AD and AR diseases. Proteins linked to AD diseases exhibited more clustering of rare missense mutations than those linked to AR diseases (Wilcoxon P=5.7×10^-4^, permutation P=8.4×10^-4^). Rare missense mutation in proteins linked to either AD or AR diseases were more clustered than controls (1000G) (Wilcoxon P=2.8×10^-15^ for AD and P=4.5×10^-4^ for AR, permutation P=3.1×10^-12^ for AD and P=0.03 for AR). Differences in clustering patterns persisted even after removal of the most prominent genes. Testing for such non-random patterns may reveal novel aspects of disease etiology in large sample studies.

## INTRODUCTION

Hermann Muller was the first geneticist to posit the existence of different classes of functional mutations effective at the protein level, mutations that he termed nullomorphs (complete loss of function), hypomorphs (reduced function), hypermorphs (increased function), antimorphs (antagonistic to wild-type) and neomorphs (new function) (1, 2). These classes of mutation can cause human disease, as well as phenotypic variability in general. Nullomorphs and hypomorphs are generally referred to today as loss-of-function mutations, and there has been speculation that they are not preferentially located at specific amino acid residue positions (2-4). This is because loss-of-function is often caused by destabilization of the hydrophobic protein core (5), or by frameshifts and premature stop codons that lead to the nonsense mediated decay (NMD) of truncated transcripts (6). On the other hand, hypermorphic, antimorphic and neomorphic mutations are generally referred to as gain-of-function mutations and are more likely to occur at specific amino acid residue positions, such as at sites of post-translational modification, ligand binding, or protein-protein interaction (5). To our knowledge, we present the first study to systematically assess and quantify the extent to which these clustering patterns are also applicable to rare missense mutations causing human inherited disease.

Single-gene diseases in which the causal mutations lie in genes residing on the autosomes are generally recognized to display either dominant (1 copy required) or recessive (2 copies) inheritance. These diseases can be caused by mutations in any of the classes mentioned above. There is a unique set of autosomal dominant diseases that are recognized to exhibit mutations in a highly restricted set of amino acid residue positions with very specific effects on protein function. By contrast, with autosomal recessive diseases, mutations are often loss-of-function and result in no or little usable protein product. Examples of specific protein functional effects include the autosomal dominant diseases Cherubism (*SH3BP2* mutations) (7) and Achondroplasia (*FGFR3* mutations) (8). In Cherubism, mutations occur at a binding site required for proper ubiquitylation and subsequent proteolytic degradation of SH3BP2(9, 10). In Achondroplasia, a mutation at residue 380 causes FGFR3 to become constitutively activated (11).

Based on the realization that mutations are often loss-of-function in recessive disease but can be either loss-of-function or gain-of-function in dominant diseases, we hypothesized that: 1) rare missense mutations within autosomal dominant (AD) disease genes might be more clustered than those in autosomal recessive (AR) disease genes; and 2) rare variants in controls might be less clustered than either. In this work, we define clustering, for a given set of mutations, as an event when mutations are closer to each other in primary protein sequence than would be expected by chance. We reasoned that if these mutation patterns generally held true, non-random clustering of rare missense mutations might provide key insights into the molecular mechanisms underlying inherited diseases. The search for new Mendelian disease genes based on whole exome sequencing is often focused on loss-of-function variants and deleterious missense variants (12). By examining non-random clustering, it becomes possible to detect regions that are critical to protein function, regardless of whether the clustered mutations are deleterious or result in gain of function.

To test the first hypothesis, we used data from The Human Gene Mutation Database (HGMD) (13), which comprises a collection of inherited mutations causing human genetic disease. To our knowledge, these data have not been previously assessed for a relationship between patterns of rare missense mutation clustering and mode of disease inheritance. To test the second hypothesis, we compared the rare missense mutations in these AD and AR genes to rare missense variants in these genes found in individuals from the 1000 Genomes Project.

First, we applied a biased approach that considered the fraction of missense mutations (or variants) in a given protein that occurred within annotated protein domains from the Human Protein Reference Database (HPRD) (14) (*domain occupancy score*). However, the assumption that rare missense mutations of large effect will only occur in protein domains, regions of regular secondary structure whose function is known and that occur paralogously in multiple proteins, is potentially problematic. Thus, we developed a new unbiased clustering method to score clustering of missense mutations in protein sequence. The method makes no *a priori* assumptions about the importance of these positions or the number of clusters.

We performed statistical testing to assess whether rare missense mutations in AD genes and AR genes exhibit different clustering patterns than in controls and from each other. AD genes were found to exhibit significantly higher protein domain occupancy than AR genes and controls, and both AD and AR genes had significantly higher occupancy than controls. When we removed the domain bias from our analysis by applying an unsupervised clustering algorithm we developed (CLUMP), we found that collectively AD genes exhibited significantly lower CLUMP scores (associated with greater clustering) than AR genes and that AD genes and AR genes had significantly lower CLUMP scores than controls. These trends persisted even after 18 outlier genes with the highest statistical significance were removed from the analysis, supporting the generality of the clustering patterns.

## RESULTS

### Generation of high quality mutations dataset and AD/AR annotations

By searching the Human Gene Mutation Database (HGMD) and using a customized pipeline (Figure 1) we generated a rare missense mutation dataset for AD genes (6,337 mutations underlying 162 diseases involving 181 genes and AR genes (6,493 mutations underlying 195 diseases involving 159 genes). A rare missense mutation was defined by a minor allele frequency < 0.01 in European controls from the 1000 Genomes Project.

**Figure 1:**
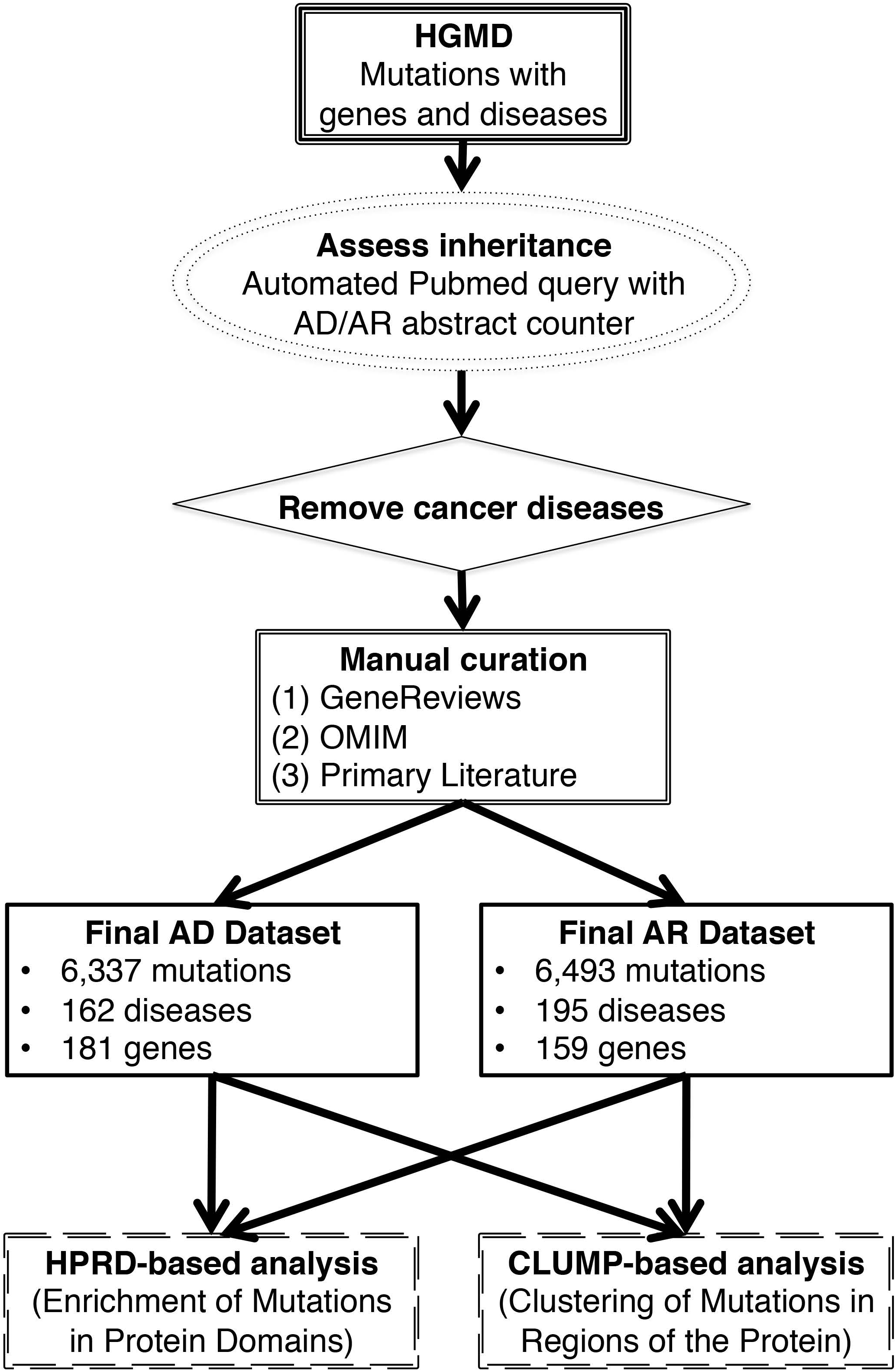
Workflow of this study. Included are details on the generation of high quality inheritance datasets for all missense variants in autosomal dominant (AD) and autosomal recessive (AR) diseases. Also depicted are our two main approaches to assess mutation clustering within proteins.

### Known disease-causing mutations are more likely to fall in domains

The general trends observed in our domain occupancy analysis are evident in (Figure 2A). The empirical cumulative distribution functions (CDFs) of domain occupancies for AD disease, AR disease, and controls (1000GP) show that the three sets are distinct and that the trend for AR disease lies midway between AD disease and controls. These trends can be further quantified by means of a non-parametric Wilcoxon test. Rare missense mutations associated with AD diseases are significantly more likely to occur within domains than are rare missense variants seen in the 1000 Genomes (p= 2.8×10^-15^, Wilcoxon test, AD median = 55%, AD mean = 55%, 1000G median = 23%, 1000G mean = 31%). Rare missense mutations associated with AR diseases also exhibit this pattern (p= 4.5×10^-4^, Wilcoxon test, AR median = 40%, AR mean = 41%) although significantly less so than those associated with AD diseases (p= 5.7×10^-4^, Wilcoxon test). In addition to these tests of mutations in individual proteins, a global analysis of all mutations shows that rare missense mutations more often reside in domains in AD diseases (total AD mutations in domains = 2,728, total AD mutations =6,337, percent AD mutations in domains = 43.0%) than in AR diseases (total AR mutations in domains = 1,771, total AR mutations=6,493, percent AR mutations in domains = 27.3%) (Fisher one-sided p=9.2×10^-79^). Generally, as previously documented (15-17) disease mutations (AD union AR) more often reside in domains than in controls (total control mutations in domains = 24,663, total control mutations=113,547, percent control mutations in domains = 21.7%) (Fisher one-sided p=6.7×10^-233^).

**Figure 2:**
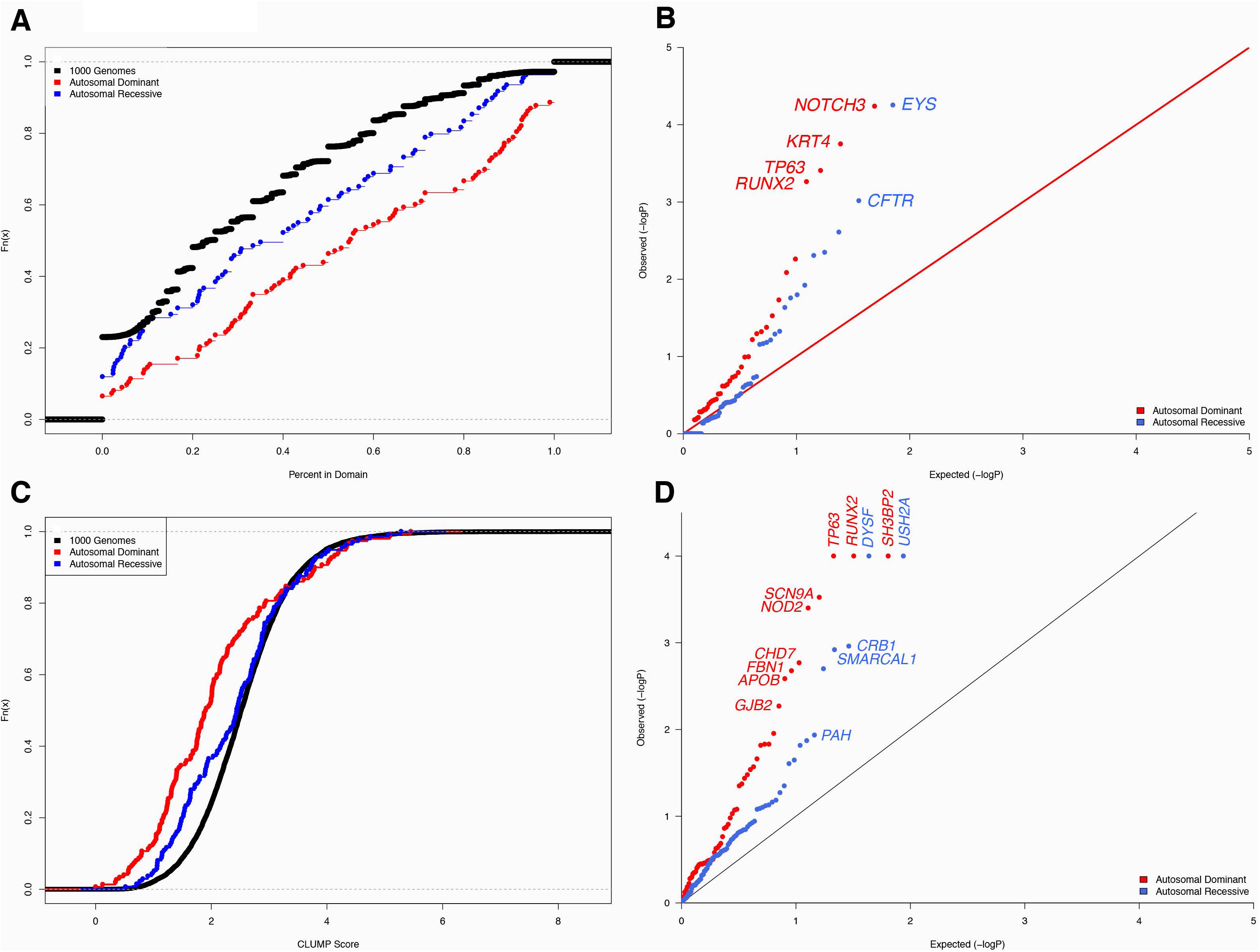
Statistical test of rare missense variant or mutation clustering within proteins. (A) Empirical cumulative distribution function of proportion of mutations residing in a domain per protein. (B) Quantile-quantile (QQ) plot of raw P-values for Fisher Exact testing to examine enrichment of mutations within domains in disease versus in controls. (C) Empirical cumulative distribution function of CLUMP scores per protein. (D) QQ plot of raw P-values for permutation testing to examine lower CLUMP scores in disease versus controls. Genes listed are those that attained a level of significance after Benjamini Hochberg correction.

**Figure 3:**
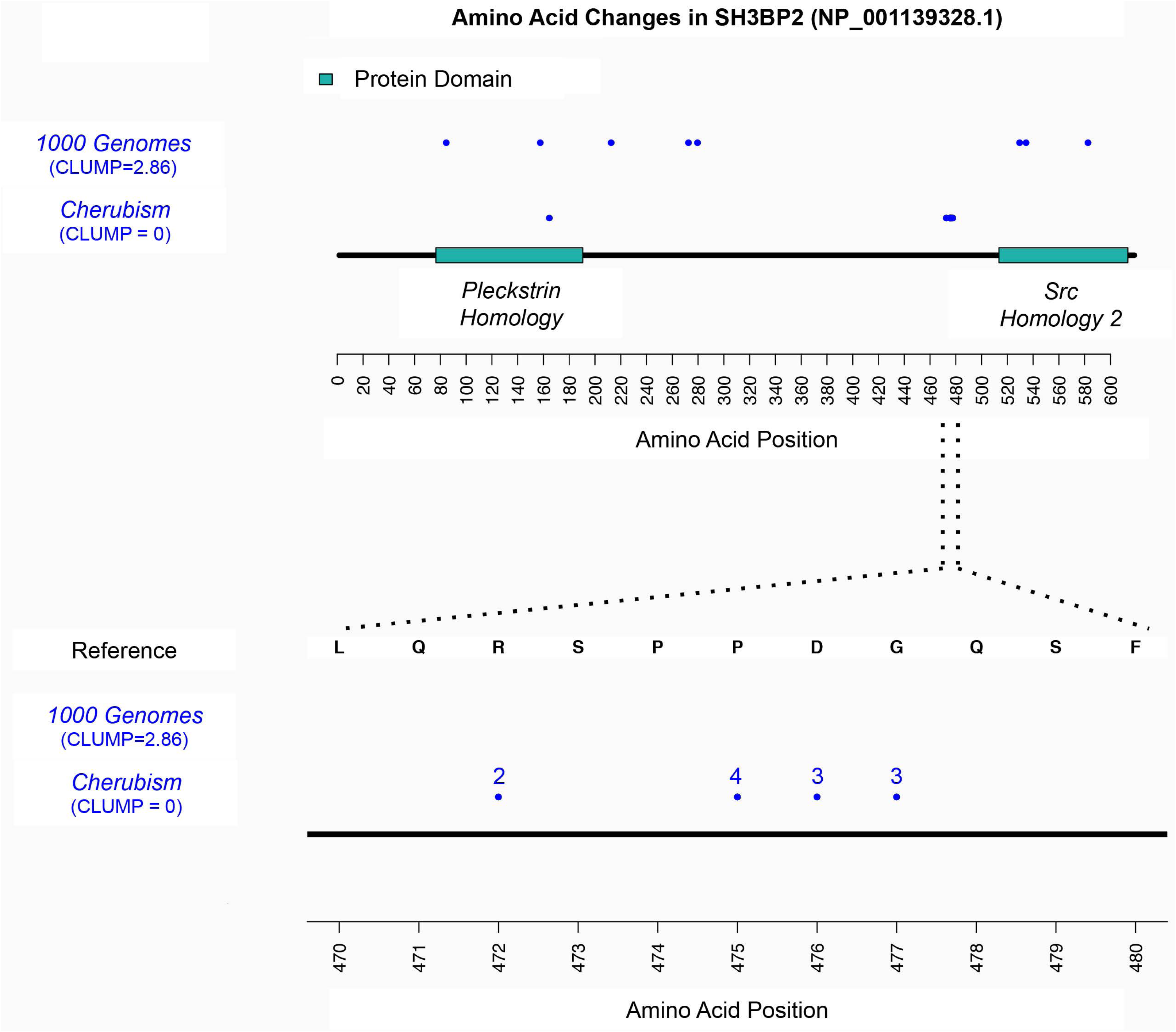
Mutations in the *SH3BP2* gene in cherubism show significant clustering. (A) Shown are all mutations in 1000 Genomes controls and in cherubism. (B) Zoom in of the region where the majority of mutations reside as well as the number of different amino acid changes at each position. The cherubism mutations are significantly more clustered than the control data (p < 1×10^-4^).

### Disease vs. control comparison of domain occupancy reveals proteins with significant differential clustering

Next, we considered whether domain occupancy could be applied to analysis of individual proteins to differentiate clustering patterns of rare missense disease mutations and control variants. We applied Fisher’s Exact test to each protein in the AD and AR sets and compared mutation clustering patterns in disease vs. controls (1000G). We identified four genes with a significant number of domain mutations in the autosomal dominant dataset and two genes in the autosomal recessive dataset, and these genes appear as outliers in a quantile-quantile (QQ) plot of raw P-values (Figure 2B). AD genes were *NOTCH3* in cerebral autosomal dominant arteriopathy with subcortical infarcts and leukoencephalopathy (CADASIL, p= 2.77×10^-3^, Benjamini-Hochberg (BH) correction), *KRT14* in epidermolysis bullosa simplex (p= 4.24×10^-3^, BH), *TP63* in ankyloblepharon-ectodermal defects-cleft lip/palate (AEC syndrome, p= 6.29×10^-3^, BH), and *RUNX2* in cleidocranial dysplasia (p=6.57×10^-3^). AR genes were *EYS* in retinitis pigmentosa (p=3.9×10^-3^, BH) and *CFTR* in cystic fibrosis (p=0.03, BH) (Figure 2B). The general trends seen in the Wilcoxon test persisted even after these outliers were removed (AD vs 1000G P=9.4×10^-14^, AR vs 1000G P=1.0×10^-3^, AD vs AR P=1.0×10^-3^)

### CLUMP analysis reveals increased clustering of autosomal dominant disease mutations

Whereas rare missense variants that occur in domains are more likely to have more influence on protein activity than those occurring outside of domains, many proteins do not have complete domain annotations (18). We further considered whether the mutation clustering trends defined by domain occupancy would persist if clustering was defined by an unbiased approach. To this end, we generated CLUMP scores for all proteins in the AD, AR and 1000 Genomes data. The empirical CDFs of CLUMP scores for AD disease, AR disease, and controls (1000G) show a similar trend to the domain occupancy scores, although the three sets are not as well separated across the full range of CLUMP scores (Figure 2C). However, the differences between the three sets remained statistically significant. Proteins with AD mutations exhibited lower scores (more clustering) than 1000 Genomes (P=3.1×10^-12^) and AR (P=8.4×10^-4^, Wilcoxon) proteins and AR proteins are themselves more localized than 1000 Genomes (P=0.03, Wilcoxon).

### Disease vs. control comparison of CLUMP scores reveals proteins with significant differential mutation clustering

To assess the statistical significance of CLUMP scores, we applied permutation testing to each protein in the AD and AR sets and compared CLUMP scores in disease vs. controls (1000G). This analysis identified 9 genes with significantly lower CLUMP scores (increased clustering) in the autosomal dominant dataset and 5 genes in the autosomal recessive dataset. Two of the AD genes were also identified in the domain occupancy analysis (*TP63* and *RUNX2*). All significant genes appear as outliers in a quantile-quantile (QQ) plot of raw P-values (Figure 2D). AD genes were *RUNX2* in cleidocranial Dysplasia, *SH3BP2* in cherubism, *TP63* in ectrodactyly, ectodermal dysplasia, clefting (EEC) syndrome, *SCN9A* in primary erythermalgia, *NOD2* in Blau syndrome, *CHD7* in CHARGE syndrome, *FBN1* in aortic aneurysm, *APOB* in hypercholesterolaemia, and *GJB2* in keratitis-ichthyosis-deafness syndrome. AR genes were *DYSF* in limb girdle muscular dystrophy, *USH2A* in Usher Syndrome, *CRB1* in Leber congenital amaurosis, *SMARCAL1* in Schimke immuno-osseous dysplasia, and *PAH* in phenylketonuria (Figure 2D). For CLUMP scores, the general trends seen in the Wilcoxon test also persisted after outliers were removed (AD vs. 1000G P = 2.5×10–^10^, AR vs. 1000G P= 0.06, AD vs. AR P = 2.3×10^-3^).

For some of these AD genes, evidence of specific protein function affected by a mutation cluster has been previously recognized. In cleidocranial Dysplasia, mutations in the transcription factor *RUNX2* cluster in the Runt domain, interfering with DNA binding (19); in EEC syndrome, mutations in the transcription factor *TP63* cluster in the DNA binding domain, disrupting DNA binding (20); and in Blau syndrome, mutations in *NOD2* cluster at its ATP-binding site and within its helical domain, dysregulating hydrolysis and autoinhibition, respectively (21).

### Proteins exhibiting increased clustering in Mendelian diseases

Of the genes whose protein products were identified to have significantly increased clustering when compared to controls, there were some that were already known to either localize in domains or cluster in a specific region of the protein. This included *RUNX2* in Clediocranial dysplasia (MIM 119600), the *TP63* gene in the AEC and EEC syndromes (MIM 603273), *SH3BP2* in Cherubism (MIM 118400), and *KRT14* in Epidermolysis bullosa simplex (MIM 148066). Our results also support the presence of a clustering pattern in the first 60 amino acid residues of the Keratitis-ichthyosis-deafness syndrome *GJB2*, which was previously observed in a small study of 10 patients (22).

### Autosomal dominant mutations are bioinformatically predicted to be more pathogenic than autosomal recessive

We have developed and published a bioinformatic variant pathogenicty classifier called the Variant Effect Scoring Tool (VEST), which outperformed SIFT or PolyPhen2 on a carefully curated benchmark set (five-fold gene holdout cross-validation cite) by a small margin (23). VEST scores range from 0 to 1 with the most having a score of 1. When we ran VEST on AD and AR variants we found that AD variants were overall more pathogenic than AR variants (Wilcoxon one-sided p=4.2×10^-10^). In addition, we found the clustered/domain variants to be more pathogenic than non-clustered/non domain variants (Wilcoxon one-sided p=3.2×10^-3^).

## DISCUSSION

A very large number of rare missense variants are now being discovered by high throughput sequencing in an assortment of human disease studies. Identifying those that are pathogenic or which contribute to disease remains very challenging. We have previously shown that visualizing the distribution of missense variants in a given protein sequence can be informative in relation to identifying potentially causal variants (24). However, such visualization does not provide quantitative assessment of clustering patterns and it cannot be applied in a high-throughput setting. In this work, we present two methods for the rapid determination of mutation clustering patterns and their statistical significance. The first method is a domain occupancy score, which considers the fraction of variants in a protein that occur within annotated domains. This score is necessarily biased, because it depends on existing knowledge of those protein regions considered to comprise functional domains, and it may miss functionally important regions that occur outside of domains. The second method is the CLUMP score, which performs unsupervised clustering of amino acid residue positions where variants occur, without any prior knowledge of their functional importance. Interestingly, we observed remarkably similar results with both methods: proteins linked to AD diseases harbor significantly more clustering of disease mutations than those linked to AR diseases, and both AD and AR disease proteins exhibit more clustering of these mutations than controls from 1000G. Moreover, these trends are not driven by a few outliers, since they persist even when the 18 genes with the most significant P-values in our Fisher’s Exact test and permutation test were removed.

It has been shown in some cases, that loss-of-function mutations (nullomorphs and hypomorphs) exhibit less clustering in protein sequence than hypermorphs and neomorphs (3, 4), but to our knowledge this is the first study to systematically assess these patterns with respect to rare missense mutations causing human inherited disease. The search for new Mendelian genes through whole exome or genome sequencing of patients has generally been focused on loss-of-function mutations (25), which have the advantage of being more readily interpretable. Bioinformatics scoring of missense mutation deleteriousness is also widespread in analysis pipelines, and features such as inter-species evolutionary conservation at a given mutation position implicitly identify amino acid substitutions that are damaging to that protein (26, 27). Often, researchers are faced with multiple rare missense variants in a gene of interest, none of which have been assessed to be damaging by popular bioinformatics tools. Our results support the idea that many of these variants may be important to Mendelian disease, but could be mutations that cause a protein gain of function and are inherited in an autosomal dominant inheritance pattern.

We have confirmed that the clustering patterns of rare missense mutations are systematically associated with mode of inheritance, and this pattern was robust with respect to whether clustering was defined by occurrence in protein domains of known functional importance or by an unbiased clustering approach. Our results are consistent with the notion that autosomal dominant disease genes harbor a mixture of deleterious and gain-of-function rare missense mutations, whereas autosomal recessive disease genes harbor only deleterious rare missense mutations.

Futher, these results suggest that sequencing studies of specific disease genes could benefit by testing for non-random clustering of rare missense variants. These clusters may provide insights into the molecular basis of inherited diseases, and such testing will become more powerful as sample sizes increase.

## MATERIALS AND METHODS

### Generation of a high quality list of disease mutations and mode of inheritance

A list of 61,537 missense mutations causing inherited disease (DM) and occurring on autosomes was downloaded from the Human Gene Mutation Database (HGMD) Professional version 2014.2 on June 10, 2014. In this study, we focused on autosomal diseases and not X-linked due to lack of information on sample sex in this dataset. For each mutation, we first parsed all abstracts in PubMed (http://www.ncbi.nlm.nih.gov/pubmed/) to identify the mode of inheritance associated with the gene in which the mutation occurred, using a custom script and BioPython libraries (28). For each entry, we generated a Boolean query of the architecture *geneName AND diseaseName AND autosomal* (example: *CFTR* AND cystic fibrosis AND autosomal). Abstracts that matched the query were then parsed for the keywords “autosomal dominant” and “autosomal recessive.” We counted the number of abstracts containing “autosomal dominant”, “autosomal recessive” or which did not contain either of these terms. An initial assignment of each entry to the autosomal dominant (AD) class, the autosomal recessive (AR) class, or as “not determined” (ND) was performed by a vote of abstracts matching these keywords, so that

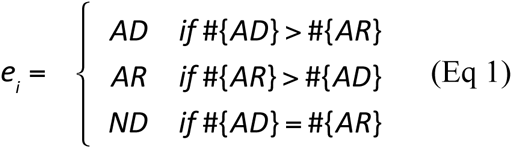

where *e*_*i*_ is an entry consisting of a gene/disease pair, #{AD} is the number of abstracts that contained the keywords “autosomal dominant”, and #{AR} is the number of abstracts that contained the keywords “autosomal recessive”. Because our study focuses on Mendelian disease, we filtered out any entries with a cancer disease association (containing the keywords cancer, sarcoma, carcinoma, leukemia, lymphoma, blastoma, glioma, melanoma, myeloma, tumor, tumour, metastasis, adenoma, neoplasia, or cytoma). At this stage, 3539 abstracts remained. To obtain high confidence calls, we further required that an entry’s classification (Eq 1) was supported by at least 12 or more abstracts and that the classification was supported by a sizeable majority (75%) of the abstracts. These criteria filtered out 80% of abstracts identified by our initial queries, yielding a high-quality set of 706 abstracts that was tractable for manual inspection. Next, every entry was manually checked for correctness of our class assignment. For each entry, we first checked for confirmation in GeneReviews (GeneTests 1999-2014), followed by OMIM (http://omim.org/), and the primary literature. Manually confirmed entries were retained.

### Control dataset

The 1000 Genomes Project dataset was obtained from ftp://ftp-trace.ncbi.nih.gov/1000genomes/ftp/ on July 18, 2014. We selected only unrelated individuals of European ancestry from the CEU, FIN, GBR, IBS, and TSI populations.

### Statistical tests for clustering of mutations and variants

To ascertain mutation clustering patterns in a gene product we adopted two approaches; the first was designed to look at the fraction of mutations occurring in annotated protein domains from the HPRD (*domain occupancy score*) and the second was the unbiased *CLUMP score*.

For a protein *p,* its *domain occupancy count* is:

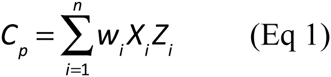

where *X*_*i*_ is a mutated amino acid residue position, *w*_*i*_ is the count of unique amino acid substitutions at that position in the data of interest, *Z*_*i*_ is binary random variable that is set to 1 when *X*_*i*_ is in an annotated protein domain, and 0 otherwise, and the sum is over the *n* mutated amino acid residue positions in the protein. Likewise,

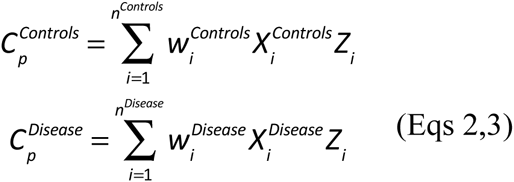

and all variables have the same meaning as in (Eq. 1) but are assigned values based only on either variants in the control set or mutations in the disease set.

For a protein *p,* its domain occupancy score (the fraction of mutations occurring in domains) is:

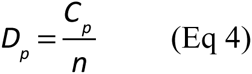

and likewise

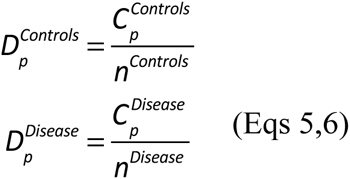

We compute 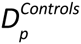 for all proteins in the control set and 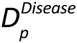 for all proteins in the disease set, and we apply a one-sided Wilcoxon test to ascertain whether the scores of proteins in the disease set are significantly higher than those in the control set. Next, to assess whether domain occupancy is significantly higher in the disease set than in the control set, for each protein we compute a one-tailed Fisher’s Exact test, comparing counts of 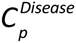, 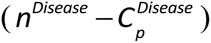, 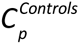, and 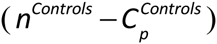. Multiple testing correction was performed with the Benjamini-Hochberg algorithm and corrected P-values < 0.05 were considered significant.

The CLUMP score applies the partitioning around medoids (PAM) clustering algorithm (29) to a list of (integer-indexed) amino acid residue positions. We use the pamk implementation in the fpc package in R. The number of clusters *k* is not specified in advance but is estimated by varying *k* over multiple PAM runs and selecting the *k\** that yields the maximum average silhouette width. Thus, both the number of clusters and a “medioid” or representative member of each cluster are estimated by the algorithm. Next, for each cluster *i*, we compute the distance between each member of the cluster and its mediod and take a log sum of these distances over all clusters. The final CLUMP score *S*_*p*_ for a protein *p* is:

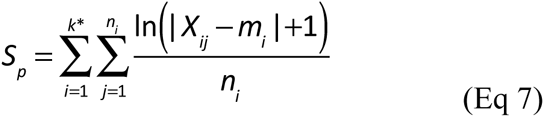

where *X*_*ij*_ is the position of mutation *j* in cluster *i*, *m*_*i*_ is the position of the mediod of cluster *i*, *n*_*i*_ is the number of mutations in cluster *i*, and *k\** is the total number of clusters in the gene. The maximum clustering possible is when all observed mutations in all clusters occur at the same position as the cluster mediod, yielding a score of 0. In general, a protein with highly localized mutations will have a low score, while a protein with mutations spread across its protein sequence will have a high score.

To assess the statistical significance of *S*_*p*_ (Eq 7), we compute for each gene’s protein product *p*, 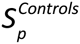 and 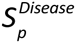 as

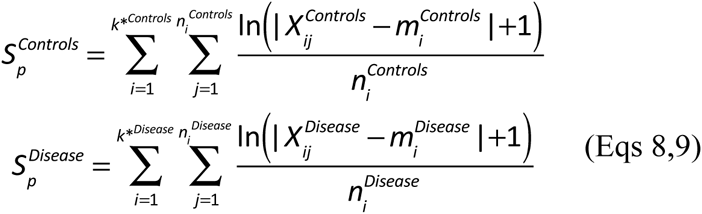

where all variables have the same meaning as in (Eq. 7) but are assigned values based only on either variants in the control set or mutations in the disease set, *i.e.,* 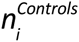 is the total number of variants observed in the protein in the control set, 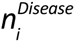 is the total number of mutations observed in the protein in the disease set, *etc.*

We compute 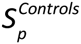 for all proteins in the control set and 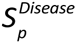 for all proteins in the disease set, and we apply a one-sided Wilcoxon test to determine if the scores of proteins in the control set are significantly higher than those in the disease set. Next, to assess whether 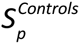 is significantly higher than 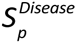 for individual proteins, we use the test statistic 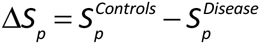.

We simulate a null distribution of values 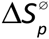 that would be expected when the difference between 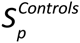 and 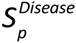 is due to random chance, by repeatedly sampling with replacement 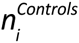 positions in protein *p* (assuming that each position is equally likely under the null hypothesis) and computing 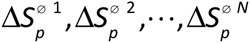, where in this work *N*=10,000. The estimated P-value for Δ*S*_*p*_ is then the fraction of times a value equal to or greater than Δ*S*_*p*_ is seen under the null. Finally we use the Benjamini-Hochberg method (30) to correct for multiple testing.

**Table 1:**
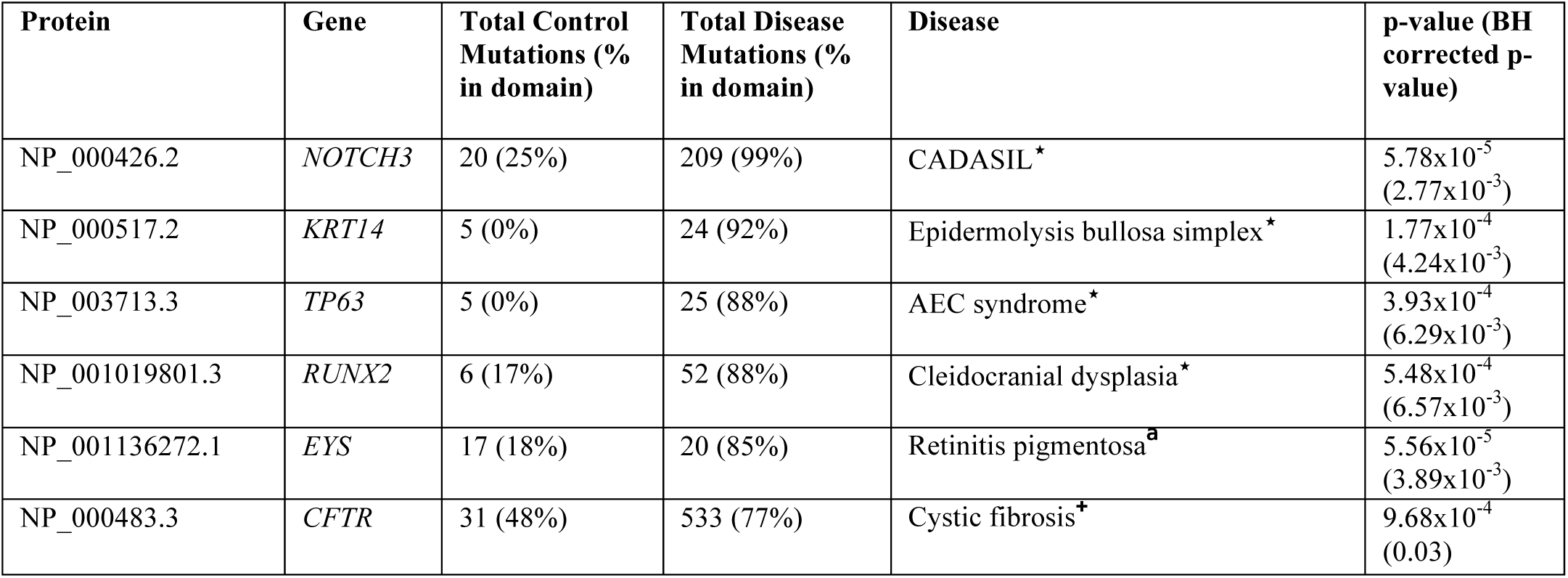
Proteins with significant enrichment of autosomal dominant and recessive rare, missense mutations in domains. Shown are counts in annotated HPRD domains or not in domains of rare (minor allele frequency < 0.01 based on controls) missense variants. The control data are from the 1000 Genomes European ancestry data. (⋆=autosomal dominant, ^**+**^=autosomal recessive).

**Table 2:**
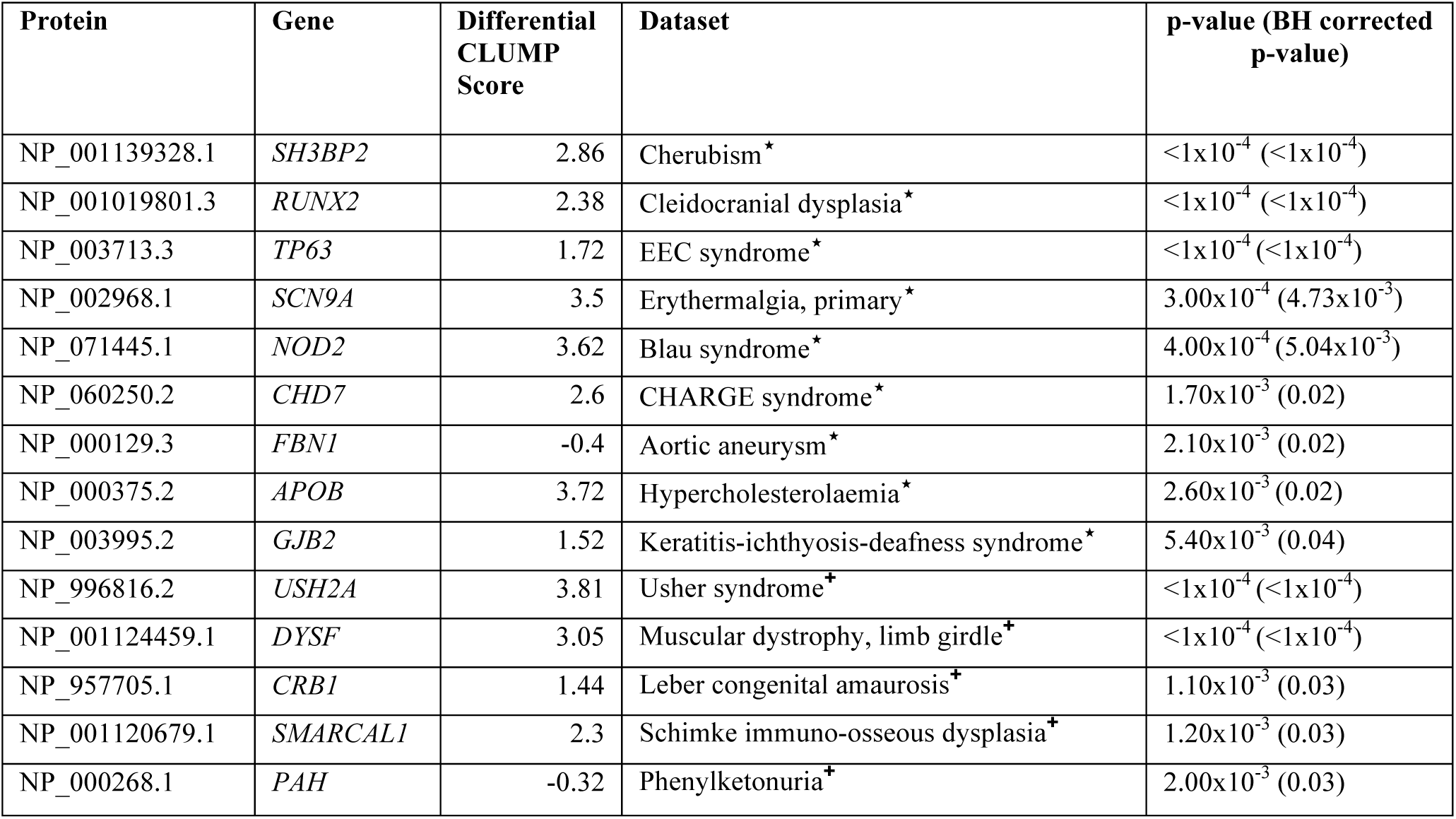
Proteins with significantly lower CLUMP scores in autosomal dominant and recessive rare, missense mutations than in controls. Shown are differential CLUMP scores between controls and disease variants of rare (minor allele frequency < 0.01 based on controls) missense variants. The control data are from the 1000 Genomes European ancestry data. (⋆ =autosomal dominant, ^**+**^=autosomal recessive).

## ACKNOWLEDGMENTS

This work was funded by a grant from the NSF (DBI-0845275) to RK. This paper began as a project in the Foundations of Computational Biology and Bioinformatics II course (Spring 2011) at the Johns Hopkins University.

### CONFLICT OF INTEREST STATEMENT

We have no conflicts of interest.

